# *β*-alanine or L-carnosine prevents the development of pale, soft and exudative pork by inhibiting the glycolysis of porcine *longissimus dorsi* muscle

**DOI:** 10.1101/2020.04.09.033225

**Authors:** Lihong Zhao, Peng Chen, Wenxiang Li, Lan Li, Yaojun Liu, Jianyun Zhang, Cheng Ji, Qiugang Ma

**Affiliations:** State Key Laboratory of Animal Nutrition, College of Animal Science and Technology, China Agricultural University, Beijing 100193, P. R. China

**Keywords:** Carnosine, *β*-alanine, L-histidine, *Longissimus dorsi* muscle, Glycolysis

## Abstract

Carnosine plays an important role in regulating muscle buffering capacity and glycolysis. In order to investigate the effects of dietary *β*-alanine, L-histidine, and L-carnosine supplementation on the pH value, glycolytic potential, the activities of AMP-activated protein kinase (AMPK) and pyruvate kinase (PK) activities in the porcine *longissimus dorsi* muscle, a total of 60 barrows with an average body weight (BW) of 50.5 ± 1.7 kg were assigned into five groups which received diets containing basal diet (control, CON), 0.04% *β*-alanine (*β*-ALA), 0.06% L-histidine (L-HIS), 0.04% *β*-alanine+0.06% L-histidine (*β*-ALA+L-HIS), or 0.1% L-carnosine (L-CAR). The results showed that dietary supplementation of the combination of *β*-ALA and L-HIS or L-CAR significantly increased (*P* < 0.05) the average daily gain (ADG) of pigs with 50-75 kg BW, compared with other three groups. Compared with L-CAR group, L-HIS supplementation significantly decreased (*P* < 0.05) the average daily feed intake (ADFI) of pigs. There were no significant difference (*P* > 0.05) in back fat thickness and loin eye area among treatments. At 0, 1 and 24 h postmortem (PM) the pH values in the *longissimus dorsi* muscle (LM) of pigs receiving L-CAR were higher (*P* < 0.05) than that of pigs receiving CON diet. The redness (a*) values of LM of pigs in the *β*-ALA+L-HIS or L-CAR group were higher (*P* < 0.05) than those in other three groups. The glycolytic potential of LM was not significantly different (*P* > 0.05) among treatments. At 1, 24 h PM the AMPK activities in LM of pigs receiving *β*-ALA, *β*-ALA+L-HIS and L-CAR were much lower (*P* < 0.05) than those of pigs receiving CON diet or L-HIS. 0.1% L-carnosine or 0.04% *β*-alanine supplemented to pigs’ diet was effective in regulating AMPK and PK activities in the porcine *longissimus dorsi* muscle and preventing the development of pale, soft and exudative (PSE) pork.

## Introduction

Carnosine (*β*-alanyl-L-histidine) is highly concentrated in skeletal muscle and brain tissues of mammals. L-histidine is an essential amino acid that can only be obtained from the diets, whereas *β*-alanine is a non-essential amino acid and the rate-limiting precursor of carnosine synthesis (1,2). The muscle carnosine contents could be augmented by oral supplementation of its two constituent amino acids. Muscle-carnosine is highly responsive to its availability in the diet with *β*-alanine supplementation (1,3).

Carnosine plays multiple biological functions including pH buffering, anti-oxidation, anti-glycation, anti-aging, and chelation of divalent metal cations (4, 5). Surface aplication of carnosine on fresh beef steaks can reduce the oxidative damages and enhance meat stability in storage by its antioxidant activity (6). Ma et al. (7) found that the dietary supplementation of carnosine did not affect the growth performance and carcass traits of pigs, but the addition of carnosine (100 mg/kg diet) increased the pH value of muscle at 45min, 24h and 48h postmortem and decreased the drip loss at 48 h postmortem. The improvement of dietary supplementation with carnosine on meat quality of pigs could be attributed to its antioxidant capacity. However, little is known about the effect of carnosine supplementation in swine diets on the postmortem muscle physiology. Renner et al. (8) studied that carnosine may interfere with glycolysis, inferring from the observations that carnosine has suppressing effects on tumor cells and differentiated fibroblasts which are predominantly glycolytic for adenosine triphosphate (ATP) supply. Degradation of glycogen through the anaerobic glycolysis is a key metabolic pathway in postmortem muscle.

Pale, soft and exudative (PSE) pork is primarily caused by a fast and excessive rate of postmortem glycolysis (9-11). In addition, AMP-activated protein kinase (AMPK) is a newly identified kinase regulating energy metabolism in vivo and has been known as the downstream target of a protein kinase cascade acting as an intracellular energy sensor, which could regulate the energy status by stimulating catabolic processes such as increasing ATP production when cells encounter an energy crisis (12,13). Therefore, the objective of this study was to investigate the effects of dietary L-carnosine, *β*-alanine and/or L-histidine supplementation on the pH value, glycolytic potential, the activities of AMPK and pyruvate kinase activities in porcine *longissimus dorsi* muscle.

## Materials and methods

### Animals and treatments

The protocol was reviewed and approved by the Animal Care and Use Committee of China Agricultural University. All procedures were performed strictly in accordance with the guidelines of the Guide for Experimental Animals of the Ministry of Science and Technology (Beijing, China), and all efforts were made to minimize suffering. L-carnosine, β-alanine and L-histidine was purchased from Shaanxi Sciphar Hi-Tech Industry Co., Ltd (Xian, China), which purity are higher than 99% according to the manufacturer.

A total of 60 purebred Yorkshire barrows with an average body weight (BW) of 50.5 ± 1.7 kg were randomly assigned into five treatment groups (12 pigs per group) which received diets containing basal diet (control, CON) 0.04% *β*-alanine (*β*-ALA), 0.06% L-histidine (L-HIS), 0.04% *β*-alanine+0.06% L-histidine (*β*-ALA+L-HIS), or 0.1% L-carnosine (L-CAR). Two pigs were housed per pen (1.8 m × 1.8 m) in an environmentally regulated finishing barn with total slatted concrete flooring. There were 6 replicate pens per treatment. Basal diets were formulated for the pigs with 50-75 kg BW and 75-100 kg BW phases. All diets were equalized on nitrogen content through supplementation of *α*-alanine (*α*-ALA). The experimental diets were formulated to meet the nutrient requirements for 50-100 kg BW pigs according to the National Research Council (2012) (Table 1).

**Table 1.** Composition and nutrient content of the basal diets (as-fed basis)

| Item | 50-75 kg BW | 75-100 kg BW |
| --- | --- | --- |
| Ingredients, % |  |  |
| Corn | 46.90 | 55.50 |
| Corn gluten meal | 5.70 | 2.40 |
| Extruded soybean | 20.00 | 16.00 |
| Soybean meal | 20.00 | 19.00 |
| Limestone | 1.20 | 1.30 |
| Dicalcium phosphate | 1.70 | 1.20 |
| Salt | 0.30 | 0.30 |
| Corn oil | 2.70 | 2.80 |
| Premix<sup>1</sup> | 1.00 | 1.00 |
| Bentonite | 0.50 | 0.50 |
| Total | 100.00 | 100.00 |
| Nutrient content, %, unless noted |  |  |
| ME, MJ/kg | 13.24 | 13.24 |
| Crude protein | 22.70 | 19.50 |
| Calcium | 0.98 | 0.88 |
| Total phosphorus | 0.64 | 0.54 |
| Available phosphorus | 0.44 | 0.35 |
| Met | 0.50 | 0.38 |
| Met + Cys | 0.90 | 0.73 |
| Lys | 1.12 | 1.00 |

<sup>1</sup>Provided per kilogram of diet: Cu, 40 mg; Fe, 80 mg; Mn, 52 mg; Zn, 140 mg; Se, 0.33 mg; I, 0.3 mg; Co, 1.4 mg; 12,040 IU of vitamin A (retinol acetate); 2,112 IU of vitamin D3; 29.7 IU of vitamin E (DL- $\alpha$ - tocopherolacetate); 2.8 mg of vitamin K3 (menadione dimethpyrimidinol); 1.2 mg of vitamin B1 (thiamin mononitrate); 7.1 mg of vitamin B2 (riboflavin); 42.9 mg of niacin; 21.6 mg of calcium pantothenate; 1.3 mg of vitamin B6 (pyridoxine); 0.12 mg of biotin; 0.44 mg of folic acid and 0.03 mg of vitamin B12; 320 mg of choline

### Feeding procedure and growth performance

Pigs were fed ad libitum with meal form feed, following a two-phase feeding model, from 50 to 75 kg BW, and from 75 to 100 kg BW. The feeding study was carried out in an automatic feeding system (FIRE, Osborne Industries, Inc., USA) in Fengze pig farm (Fujian, China), and the time, duration and feed consumption are recorded for each time when a pig visits the feeder (*ad libitum* feeding). The initial and final body weight and feed intake of pigs of each replicate pen were recorded to calculate the growth performance including average daily gain (ADG), average daily feed intake (ADFI) and feed to gain ratio (F/G) in each feeding stage.

### Muscle samples collection and carcass measurements

When the mean weight of pigs reached around 100 kg, all pigs was transported to slaughterhouse 8 km away. Feed was withheld for 24 h before slaughtering and the pigs were laired for 4 h with free access to water before being electronically stuned, then slaughtered at the same day. After the pigs were exsanguinated, back fat thickness (measured with an A-mode ultrasound, Lean-Meater, Renco Corporation, Minneapolis, USA) and loin eye area at the 10th rib were measured. Back fat thickness was the average of measurements at three points: the first rib, the last rib, and the last *lumbar vertebra*, depth, as described by D’Astous-Pagé et al.(2017a).

The *longissimus dorsi* muscles (LM) were removed between *thoracic vertebrae* 10-13 in the left side of pig carcasses, vacuum packed, and stored after dividing into two parts (one for measurement of meat quality traits at 4°C and another for glycolytic potential and enzyme activities analysis at −80°C).

### Measurement of meat quality traits

A portable pH meter (HI99161, Hanna Instruments Inc., Italy) equipped with a spear-tipped glass pH electrode was used for pH measurement of chops of LM at 0 h PM. After the determination of LM pH at 0 h PM, the chops of LM were vacuum-packed at 4 °C to measure subsequently meat quality, including pH at 1, 24 h PM, and meat color. The meat color was assessed using a Minolta chromameter (CR-300, Minolta Camera, Japan). The average of triplicate measurements was recorded, and the results were expressed as C.I.E. (Commission Internationale de l’Eclairage) lightness (L*), redness (a*), and yellowness (b*) values.

### Detection of glycolytic potential and enzyme activities in LM

For the glycolytic potential, 1 g of frozen longissimus thoracis obtained was homogenized with 10 ml of 0.55 mol/L perchloric acid for the determination of glycogen, glucose, glucose-6-phosphate and lactate according to the method of Monin and Sellier (1985), and glycolytic potential (GP; reported in mol/g of wet tissue) was calculated according to the formula: GP = 2 × [(glycogen) + (glucose) + (glucose-6-phosphate)] + (lactic acid) (Monin and Sellier 1985). The AMP-activated protein kinase (AMPK) and pyruvate kinase (PK) activities in the LM were determined by assay kits produced by Nanjing Jiancheng Bioengineering Institute (Nanjing, China).

### Statistical analyses

Data were analyzed by one-way analyses of variance (ANOVA) using the Proc General Linear Model (GLM) procedures of the SAS statistical package (V8.1, SAS Institute Inc., Cary, NC, USA), and the statistical model is Yij = μ + αi + eij (i = 1, 2, 3, 4, 5; j = 1 to 6), where Yij is the observation of the jth animal in the ith treatment, μ is the overall mean, αi is the effect of treatment, eij is the random error and the variance between measures within pigs. Each animal served as the experimental unit. Difference among means was tested by the Student-Newman-Keuls (SNK) method. The significance level is *P* ≤0.05.

## Results

### Growth performance and carcass traits

Dietary supplementation of the combination of *β*-ALA and L-HIS or L-CAR significantly increased (*P* < 0.05) the ADG of pigs from 50 to 75 kg BW, compared with the other three groups. At the phases of 50-75 kg and 75-100 kg BW, pigs in L-CAR group had higher (*P* < 0.05) ADFI than those in L-HIS group. However, there were no significant differences (*P* > 0.05) among CON, *β*-ALA and *β*-ALA+L-HIS groups. Besides, no significant differences (*P* > 0.05) were found in F/G from 75 to 100 kg BW, and fat thickness and loin eye area between treatments.

### Meat quality

Pigs receiving L-CAR had higher (*P* < 0.05) pH values in LM than those fed with CON or L-HIS diet and β-ALA was significant lower than L-CAR at 1h and 24 h, however no differences (*P* > 0.05) were found between L-CAR and *β*-ALA+L-HIS group at 0, 1, 24 h PM (Table 3). Moreover, there were no differences (*P* > 0.05) in pH 0, 1, 24 h PM values of LM among CON, *β*-ALA and *β*-ALA+L-HIS groups. The a* values of LM in the *β*-ALA+L-HIS or L-CAR group were higher (*P* < 0.05) than those in other three groups. There were no differences (*P* > 0.05) in L* and b* values among the groups (Table 3).

**Table 2.** Effects of dietary *β*-alanine, L-histidine, and L-carnosine on the growth performance and carcass traits of pigs from 50 to 100 kg body weight (BW)

| Item<sup>1</sup> | CON | β-ALA | L-HIS | β-ALA<br>+L-HIS | L-CAR | SEM | *P* value |
| --- | --- | --- | --- | --- | --- | --- | --- |
| 50-75 kg BW |  |  |  |  |  |  |  |
| ADG, g | 0.773<sup>b</sup> | 0.780<sup>b</sup> | 0.769<sup>b</sup> | 0.862<sup>a</sup> | 0.875<sup>a</sup> | 0.02 | 0.015 |
| ADFI, g | 2.86<sup>ab</sup> | 2.72<sup>ab</sup> | 2.49<sup>b</sup> | 2.57<sup>ab</sup> | 2.96<sup>a</sup> | 0.14 | 0.036 |
| F/G, g/g | 3.71 | 3.48 | 3.23 | 2.99 | 3.36 | 0.32 | 0.078 |
| 75-100 kg BW |  |  |  |  |  |  |  |
| ADG, g | 0.847<sup>ab</sup> | 0.866<sup>ab</sup> | 0.833<sup>b</sup> | 0.868<sup>ab</sup> | 0.948<sup>a</sup> | 0.02 | 0.023 |
| ADFI, g | 3.11<sup>ab</sup> | 2.98<sup>ab</sup> | 2.75<sup>b</sup> | 2.82<sup>ab</sup> | 3.20<sup>a</sup> | 0.06 | 0.031 |
| F/G, g/g | 3.67 | 3.44 | 3.32 | 3.23 | 3.37 | 0.25 | 0.18 |
| Back fat thickness, cm | 2.39 | 2.55 | 2.24 | 2.69 | 2.74 | 0.32 | 0.22 |
| Loin eye area, cm<sup>2</sup> | 38.7 | 39.4 | 38.7 | 40.7 | 41.8 | 2.34 | 0.32 |

<sup>1</sup>CON, the control basal diet; β-ALA, the basal diet supplemented with 0.04% β-alanine; L-HIS, the basal diet supplemented with 0.06% L-histidine; β-ALA+L-HIS, the basal diet supplemented with 0.04% β-alanine and 0.06% L-histidine; L-CAR, the basal diet supplemented with 0.10% L-carnosine; SEM, standard error of mean; ADG, average daily gain; ADFI, average daily feed intake; F/G, feed to gain ratio
<sup>a,b</sup>Within a same row, means with different superscripts differ significantly ( $P < 0.05$ )

**Table 3.** Effects of dietary *β*-alanine, L-histidine, and L-carnosine on the meat quality of the *longissimus dorsi* muscle in pigs (n=6)

| Item<sup>1</sup> | CON | β-ALA | L-HIS | β-ALA<br>+L-HIS | L-CAR | SEM | P value |
| --- | --- | --- | --- | --- | --- | --- | --- |
| Meat color |  |  |  |  |  |  |  |
| Lightness (L*) | 46.2 | 46.0 | 46.3 | 45.2 | 43.1 | 1.01 | 0.47 |
| Redness (a*) | 15.4<sup>b</sup> | 15.9<sup>b</sup> | 15.3<sup>b</sup> | 17.5<sup>a</sup> | 17.8<sup>a</sup> | 1.08 | 0.016 |
| Yellowness (b*) | 5.44 | 5.65 | 5.54 | 5.60 | 5.62 | 0.12 | 0.58 |
| pH values postmortem (PM) |  |  |  |  |  |  |  |
| 0 h PM | 6.36<sup>b</sup> | 6.42<sup>ab</sup> | 6.32<sup>b</sup> | 6.62<sup>a</sup> | 6.68<sup>a</sup> | 0.05 | 0.015 |
| 1 h PM | 5.77<sup>b</sup> | 5.79<sup>b</sup> | 5.75<sup>b</sup> | 5.84<sup>ab</sup> | 6.12<sup>a</sup> | 0.10 | 0.028 |
| 24 h PM | 5.56<sup>b</sup> | 5.58<sup>b</sup> | 5.52<sup>b</sup> | 5.60<sup>ab</sup> | 6.03<sup>a</sup> | 0.02 | 0.021 |

<sup>1</sup>CON, the control basal diet; β-ALA, the basal diet supplemented with 0.04% β-alanine; L-HIS, the basal diet supplemented with 0.06% L-histidine; β-ALA+L-HIS, the basal diet supplemented with 0.04% β-alanine and 0.06% L-histidine; L-CAR, the basal diet supplemented with 0.10% L-carnosine; SEM, standard error of mean;
<sup>a,b</sup>Within a same row, means with different superscripts differ significantly ( $P < 0.05$ )

**Table 4.** Effects of dietary *β*-alanine, L-histidine, and L-carnosine on the glycolytic potential and enzyme activities of the *longissimus dorsi* muscle in pigs (n=6)

| Item<sup>1</sup> | CON | β-ALA | L-HIS | β-ALA<br>+L-HIS | L-CAR | SEM | P value |
| --- | --- | --- | --- | --- | --- | --- | --- |
| 0 h postmortem |  |  |  |  |  |  |  |
| GP,<br>$\mu\text{mol/g}$ | 153 | 150 | 151 | 130 | 124 | 3.32 | 0.28 |
| AMPK,<br>$\text{nmol/peptide/min/g}$ | 1.25 | 1.24 | 1.29 | 1.28 | 1.30 | 0.03 | 0.23 |
| PK,<br>$\text{U/ mg pro}$ | 12.4 | 12.3 | 12.4 | 12.2 | 12.1 | 0.15 | 0.29 |
| 1 h postmortem |  |  |  |  |  |  |  |
| GP,<br>$\mu\text{mol/g}$ | 160 | 156 | 155 | 132 | 130 | 2.25 | 0.21 |
| AMPK,<br>$\text{nmol/peptide/min/g}$ | 3.76<sup>a</sup> | 2.95<sup>b</sup> | 3.64<sup>a</sup> | 2.82<sup>b</sup> | 2.77<sup>b</sup> | 0.22 | 0.032 |
| PK,<br>$\text{U/ mg pro}$ | 12.0 | 11.8 | 11.8 | 11.3 | 11.3 | 0.23 | 0.17 |
| 24 h postmortem |  |  |  |  |  |  |  |
| GP,<br>$\mu\text{mol/g}$ | 177 | 168 | 176 | 151 | 138 | 3.61 | 0.25 |
| AMPK,<br>$\text{nmol/peptide/min/g}$ | 2.80<sup>a</sup> | 2.24<sup>b</sup> | 2.84<sup>a</sup> | 2.03<sup>b</sup> | 1.95<sup>b</sup> | 0.18 | 0.035 |
| PK,<br>$\text{U/ mg pro}$ | 8.56<sup>a</sup> | 8.24<sup>a</sup> | 8.31<sup>a</sup> | 7.42<sup>b</sup> | 7.28<sup>b</sup> | 0.26 | 0.027 |

<sup>1</sup>CON, the control basal diet; β-ALA, the basal diet supplemented with 0.04% β-alanine; L-HIS, the basal diet supplemented with 0.06% L-histidine; β-ALA+L-HIS, the basal diet supplemented with 0.04% β-alanine and 0.06% L-histidine; L-CAR, the basal diet supplemented with 0.10% L-carnosine; SEM, standard error of mean;
<sup>a,b</sup>Within a same row, means with different superscripts differ significantly ( $P < 0.05$ )

### Glycolytic potential enzyme activities in LM

There were no differences (*P* > 0.05) in glycolytic potential, and AMPK, PK activities of LM among treatments at 0 h PM and there is no effect of treatment on PK activity at 1 h post-mortem. Pigs fed with *β*-ALA, *β*-ALA+L-HIS or L-CAR diet had lower (*P* < 0.05) AMPK activities in LM than pigs receiving CON or L-HIS diet, and no differences (*P* > 0.05) were found in glycolytic potential and AMPK activities in LM among *β*-ALA, *β*-ALA+L-HIS and L-CAR groups at 1, 24 h PM. Pigs in *β*-ALA+L-HIS or L-CAR group had lower PK activities (*P* < 0.05) in LM than those fed with CON, *β*-ALA and L-HIS diets, and no differences (*P* > 0.05) were found among the CON, *β*-ALA and L-HIS groups at 24 h PM.

## Discussion

Diet has been shown to influence the concentration of carnosine in muscle. Like In many cases, the role of *β*-alanine as a precursor of carnosine has been found to improve the concentration of carnosine in muscle (1, 14). Dietary *β*-alanine plus L-histidine supplementation increased carnosine concentrations of type IIA and IIB muscle fibers in horses (3). In humans, dietary supplementation of *β*-alanine increased carnosine concentrations of the soleus and gastrocnemius muscles (15).

The present result indicated that dietary addition of L-CAR improved the ADG of pigs at the body weight of 50-75 kg, which was consistent with the result of Bao et al. (16) showing dietary supplementation with carnosine presented beneficial effect on growth performance of finishing pigs. In addition, dietary L-histidine supplementation significantly decreased the ADFI of pigs compared with the L-carnosine supplementation. A similar result was also reported by Haug et al. (17) that the feed intake of broilers at d 28 was lower with high histidine supplementation in diet. Some researches have reported that histidine have a suppressive effect on food intake (18,19). Both intraperitoneal injection of L-HIS and dietary supplementation of L-HIS suppressed food intake by means of its conversion into histamine in animal’s hypothalamus (20, 21). It is noteworthy to mention that the combination of *β*-alanine (*β*-ALA) and L-HIS also increased the ADG of pigs at the body weight of 50-75 kg, however, the ADFI of pigs in this group was not decreased as much as in L-HIS group. These results suggest that supplementation with *β*-ALA plus L-HIS may be more effective in increasing carnosine levels in muscle than supplementation with *β*-ALA alone, and even to some extent compensate the negative effect of L-HIS on feed intake, but the possible mechanisms need to be further elucidated.

The pH of muscle will be changed after slaughter, which due to the degradation of glycogen to lactic acid by glycogenolysis and glycolysis under anaerobic conditions. The accumulation of lactic acid reduce the pH in the muscle during enhanced anaerobic glycolysis. Notably, a rapid pH drop in muscle during the first 60 min postmortem can lead to the PSE pork production (22). To improve the meat quality of pigs, several potential inhibitors of the glycolysis have been supplemented in the diet to prevent or reduce the pH drop in muscle. It has been believed that histidine and its related peptides carnosine (*β*-alanyl-L-histidine) are components as muscle buffers, and the pH-lowering effect of lactate production may also be influenced by an increase in the buffering capacity of porcine postmortem muscle (23).

Results of the present study demonstrate that carnosine is effective in maintaining higher pH value of muscle. It has been suggested that high carnosine levels in glycolytic tissue are needed to maintain stable pH by buffering the big amounts of protons produced as a result of glycolytic activity such as through lactic acid formation and to combat the potentially deleterious by-products of glycolysis such as methylglyoxal (24, 25), the specific correlation between carnosine concentrations and metabolic state is not clear. Moreover, our study showed that dietary L-HIS didn’t increase the pH value of *longissimus dorsi* muscle, which is consistent with the result of Haug et al. (17) showing dietary histidine supplementation did not affect pH in broilers breast muscle 6 h postmortem. However, Forde-Skjaervik et al. (26) reported that the feed containing 48 g histidine/kg feed gave a significantly higher content of free histidine, higher muscle pH in white muscle of farmed atlantic cod (*Gadus morhua*). In the present study there was no effect of treatments on backfat thickness and loin eye area.

In order to reflect the ability of compounds in the muscles to convert to lactate, the muscle metabolites glucose, glucose 6-phosphate, glycogen and lactate has been be combined into a single measure termed glycolytic potential (GP), indicating the muscle’s capacity for extended glycolysis postmortem (27). In the current study, muscle glycolytic potential was measured in the LM containing fast-twitch glycolytic fibers predominantly, and the results suggested that no difference in GP was found among treatments at 0 h postmortem. This result can be partially elucidated by the fact that the glycolytic capacity of the porcine muscles increased with the increase of quantities of carnosine or *β*-alanine (28). High levels of glycolytic potential could only explain roughly 35 to 50% of the low pH and PSE-like characteristics in muscles (29). In theory, the duration and termination of glycolysis are regulated by substrate availability and enzyme activity in muscles postmortem (9). It might explain that the pH values were different among treatments in the current study, although there were not significant differences in glycolytic potential.

It is becoming increasingly recognized that AMPK activation are involved in sensing and responding to cell stress, including ischemic cardiac and skeletal muscle (30-32). Similarly, when the animals were slaughtered, most of muscle fibers will become hypoxic-ischemic and the postmortem energy metabolism of muscles converts glycogen into lactic acid by means of the glycolytic pathway. The involvement of AMPK in modulating postmortem muscle metabolism was confirmed in several studies on mice (33-35) and pigs (11, 36), which was firstly evidenced in pig muscle through the discovery of a mutation of the AMPK*γ*3 subunit responsible for a 70% increase in muscle glycogen and the production of low ultimate pH, reduced water-holding capacity, and poor processing ability meat (37, 38). These studies have demonstrated that AMPK activation can affect both the early rate and the extent of pH decrease in postmortem muscle, therefore it is a potential molecular target for the regulation of meat quality (39).

In this study, AMPK was activated in postmortem LM from pigs in *β*-ALA, *β*-ALA+L-HIS and L-CAR group, with the higher activity at 1 h postmortem. The sustained activation of AMPK in muscle should be due to the fact that ATP is rapidly consumed and a high level of AMP accumulates during postmortem change (33). No literature report the effect of carnosine on AMPK activity in muscleat present. Results of this study demonstrated that feeding carnosine decreased AMPK activity at 1 and 24 h postmortem. Additionally, pigs fed with *β*-ALA or *β*-ALA+L-HIS also had lower AMPK activity at the same stages. In this sense, we could speculate dietary supplementation of *β*-alanine increased the concentration of muscle carnosine, and the latter one inhibits the initiation and progress of glycolysis by inhibiting AMPK activation in porcine muscle postmortem.

Pyruvate kinase (PK) is a key glycolytic enzyme which irreversibly catalyzes the conversion of phosphoenolpyruvic acid to pyruvic acid and has been found to exhibit fourfold higher activity in PSE than in normal muscles. The phosphorylation of PK results in the increase of enzyme activity in PSE meat (10). Shen and Du (33) suggested that AMPK activity can influence the activity of PK, which is in agreement with our results showing that significant difference in PK activity was detected among treatments at 24 h postmortem. This effect may be due to the fact that carnosine decreases the PK activity by suppressing AMPK activity, and then reduces production of pyruvate by glycolysis, but the possible mechanisms need to be further elucidated. Taken together, dietary *β*-ALA or carnosine could reduce the activation of AMPK in the porcine *longissimus dorsi* muscle. Carnosine or *β*-alanine prevents the development of pale, soft and exudative pork by inhibiting the glycolysis of porcine *longissimus dorsi* muscle.

## CRediT author statement

**Lihong Zhao:**Methodology, Formal analysis, Validation, Writing-Original Draft, Writing-Review & Editing. **Peng Chen:** Investigation, Formal analysis, Validation, Methodology, Writing-Original Draft. **Qiugang Ma:** Conceptualization, Methodology, Project administration, Funding acquisition, Supervision. **Cheng Ji:** Supervision, Methodology. **Wenxiang Li:** Writing-Original Draft. **Lan Li:** Writing-Review & Editing. **Yaojun Liu:** Writing-Original Draft. **Jianyun Zhang:** Supervision, Writing - Review & Editing.

## Acknowledgements

We greatly appreciate the supports of the National Natural Science Foundation of China (Grant No.31572447), China Program for New Century Excellent Talents in University (NCET-13-0558) and a Sino German cooperative fund between China Agricultural University(CAU) and Hohenheim University(UOH).

## Compliance with ethical standards

Informed consent was obtained from all individual participants included in the study.

## Ethical standards

The protocol was reviewed and approved by the Animal Care and Use Committee of China Agricultural University.

